# Lower complexity of motor primitives ensures robust control of high-speed human locomotion

**DOI:** 10.1101/2020.04.24.055277

**Authors:** Alessandro Santuz, Antonis Ekizos, Yoko Kunimasa, Kota Kijima, Masaki Ishikawa, Adamantios Arampatzis

## Abstract

Walking and running are mechanically and energetically different locomotion modes. For selecting one or another, speed is a parameter of paramount importance. Yet, both are likely controlled by similar low-dimensional neuronal networks that reflect in patterned muscle activations called muscle synergies. Here, we investigated how humans synergistically activate muscles during locomotion at different submaximal and maximal speeds. We analysed the duration and complexity (or irregularity) over time of motor primitives, the temporal components of muscle synergies. We found that the challenge imposed by controlling high-speed locomotion forces the central nervous system to produce muscle activation patterns that are wider and less complex relative to the duration of the gait cycle. The motor modules, or time-independent coefficients, were redistributed as locomotion speed changed. These outcomes show that robust locomotion control at challenging speeds is achieved by modulating the relative contribution of muscle activations and producing less complex and wider control signals, whereas slow speeds allow for more irregular control.

## Introduction

Humans can locomote at a very broad range of speeds even though walking and running, the two most common gait modes, are profoundly different from both a mechanic and energetic point of view [1,2]. Walking, with its characteristic double support stance phase, typically implies at least one limb being in contact with the ground, while running allows for a flight phase [3]. Moreover, the energy cost function of walking has a peculiar U-shape, with a minimum close to each individual’s preferred speed, which lies around 1.4 m/s in the average human [4]. At lower and higher speeds, walking is relatively costlier but remains more economical than running until circa 2.4 m/s, speed at which running becomes more economical than walking [1,4]. Humans often decide to switch from walking to running at lower speeds [4], on average around 2.0 m/s. The cost of running is quasi-linearly correlated with speed, at least if the nonlinear contribution of air resistance is neglected [4–9]. Yet, despite the profound mechanical and energetic differences, walking and running seem to be sharing similar neural control [3,10,11].

The exceptional amount of degrees of freedom available to vertebrates for accomplishing any kind of movement is defined by the vast number of muscles and joints [12]. Nevertheless, the central nervous system (CNS) manages to overcome complexity, possibly through the orchestrated activation of functionally-related muscle groups, rather than through muscle-specific commands [12,13]. These common activation patterns, called muscle synergies, might be a means for the CNS to simplify the motor control problem by reducing its dimensionality [14]. Usually extracted from electromyographic (EMG) data via linear machine learning approaches such as the non-negative matrix factorisation (NMF), muscle synergies have been increasingly employed in the past two decades for providing indirect evidence of a simplified, modular control of movement in humans and other vertebrates [15–19].

In this study, we extracted muscle synergies from the EMG activities of lower limbs during treadmill walking and running at several speeds, from slow walking to maximal sprinting. Synergies were divided into time-independent (motor modules) and time-dependent (motor primitives) coefficients. The Higuchi’s fractal dimension (HFD) was used to evaluate the complexity of motor primitives, taken as self-affine time series [11,20,21]. Defining robustness as the ability to cope with perturbations [10], it follows that biological systems can manage to maintain function despite disturbances only through robust control [22–24]. Assessing the complexity of control signals could give us an idea of the strategies adopted by the CNS to cope with disruptions. Recently, we showed that challenging locomotion conditions (i.e. in the presence of external mechanical perturbations and in aging) manifest lowered complexity of motor primitives [11]. Moreover, we and others proposed that the width of motor primitives increases to ensure robust control in the presence of internal and external perturbations [10,19,25–27], suggesting that this might be a compensatory mechanism adopted by the CNS to cope with the postural instability of locomotion in health and pathology [25,26]. We observed the neural strategy of motor primitive widening in wild-type mice [19] and in humans affected by multiple sclerosis [28] or healthy adults undergoing external perturbations [10,11], but not in genetically modified mice that lacked feedback from proprioceptors [19]. Due to these observations, we concluded that intact systems use wider (i.e. of longer duration) and less complex control signals to create an overlap between chronologically-adjacent synergies to regulate motor function through robust control [10,11,28].

Here, we used the challenges imposed by slow and increasingly high speeds to perturb the locomotor system. We hypothesized that forcing the CNS to control increasingly higher speeds would perturb the system to the point of eliciting an increased control’s robustness. We discovered that motor primitives become wider and less complex as locomotion speed increases, translating into robust control. Moreover, we found that walking and running shared similar motor modules that were regulated depending on speed, confirming previous results obtained by other authors [3,29–32]. These findings provide further insight into how the CNS might control challenging locomotion. A topic with broad implications in human pathology and performance, robotics, comparative biology and other locomotion-related fields.

## Results

### Muscle synergies

The EMG activities from which muscle synergies were extracted are presented in Figure 1 as average of all trials. The average number of synergies which best accounted for the EMG data variance (i.e. the factorisation rank) of G1 was 4.3 ± 0.6 (walking, 0.7 m/s), 4.4 ± 0.5 (walking, 1.4 m/s), 4.3 ± 0.5 (walking, 2.0 m/s), 4.1 ± 0.4 (running, 2.0 m/s), 4.3 ± 0.7 (running, 3.0 m/s), and 4.3 ± 0.6 (running, 3.5 m/s). In G2, the values were 4.1 ± 0.5 (running, 2.8 m/s), 3.9 ± 0.6 (running, 4.2 m/s), 3.9 ± 0.5 (running, 5.6 m/s), 3.9 ± 0.5 (running, 6.9 m/s), 4.0 ± 0.0 (running, 8.3 m/s), and 4.2 ± 0.4 (running, 9.5 m/s). We did not find a significant effect of speed on the factorisation rank (p = 0.797 for G1, p = 0.320 for G2).

**Figure 1.**
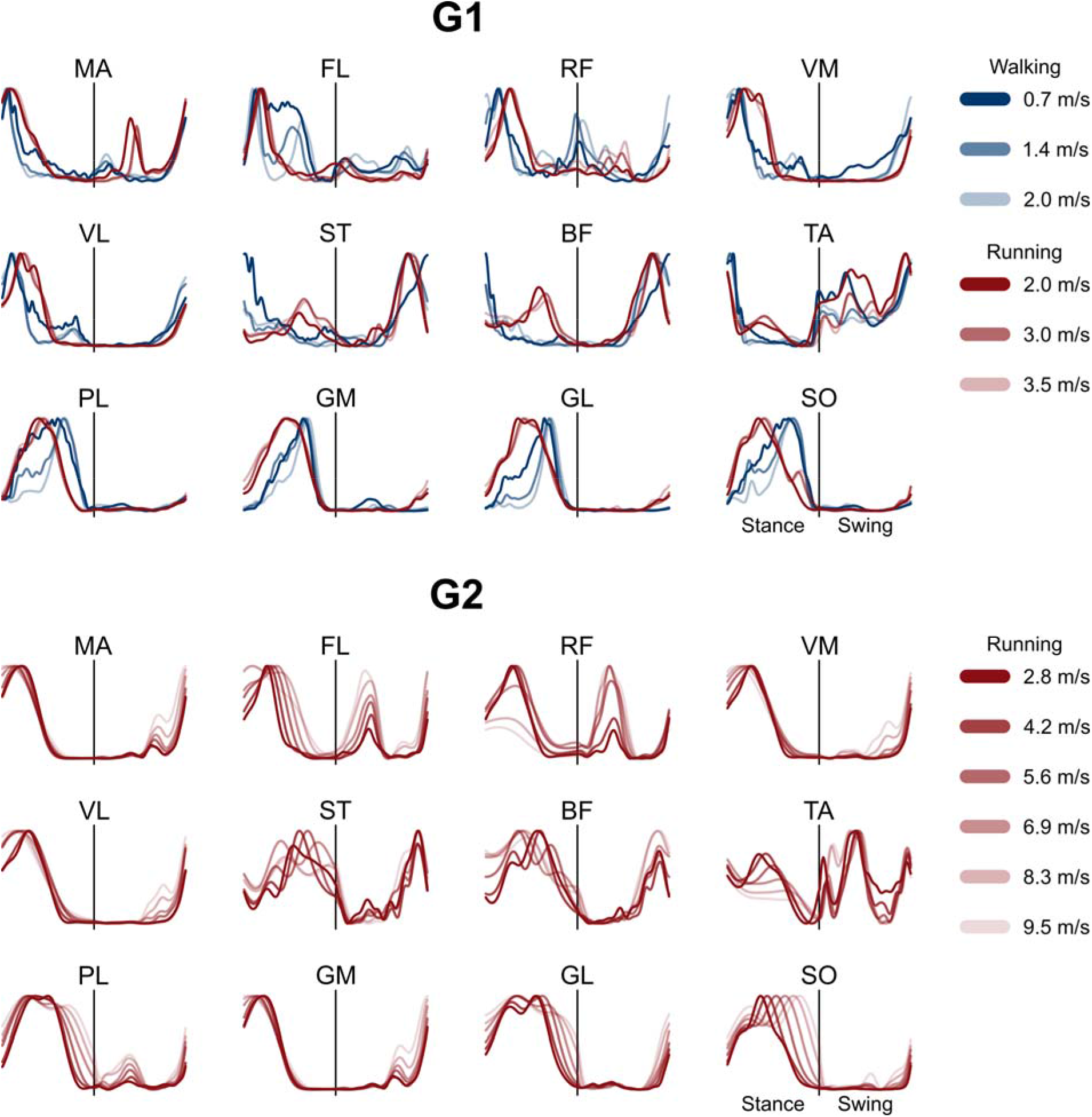
Average electromyographic activity of lower limb muscles. Average EMG activity of the recorded muscles at different speeds in group 1 (G1) and group 2 (G2). The x-axis full scale represents the averaged gait cycle (with stance and swing normalized to the same amount of points and divided by a vertical line) and the y-axis the amplitude normalized to the maximum. Muscle abbreviations: MA=*gluteus maximus*, FL=*tensor fasciæ latæ*, RF=*rectus femoris*, VM=*vastus medialis*, VL=*vastus lateralis*, ST=*semitendinosus*, BF=*biceps femoris*, TA=*tibialis anterior*, PL=*peroneus longus*, GM=*gastrocnemius medialis*, GL=*gastrocnemius lateralis*, SO=*soleus*.

The functional classification (see methods) identified four fundamental muscle synergies in both groups (Figure 2 and Figure 3). The first synergy functionally referred to the body weight acceptance, with a major involvement of knee and hip extensors. The second synergy was associated with the propulsion phase, to which the plantarflexors mainly contributed. The third synergy identified the early swing and showed the involvement of foot dorsiflexors and, at high locomotion speeds in both walking and running, of hip flexors. The fourth and last synergy reflected the late swing and the landing preparation, highlighting the relevant influence of knee flexors and foot dorsiflexors. As showed in the past for other locomotion conditions [10,33,35,38,52], not all the participants exhibited all the four fundamental synergies at all speeds; in particular, 27% and 30% of the total synergies were classified as combined in walking and running, respectively. We reported the detailed numbers in Table 1. The effect of speed on motor modules is reported in Figure 2 and Figure 3, where asterisks denote the outcome of the *post-hoc* analysis.

**Figure 2.**
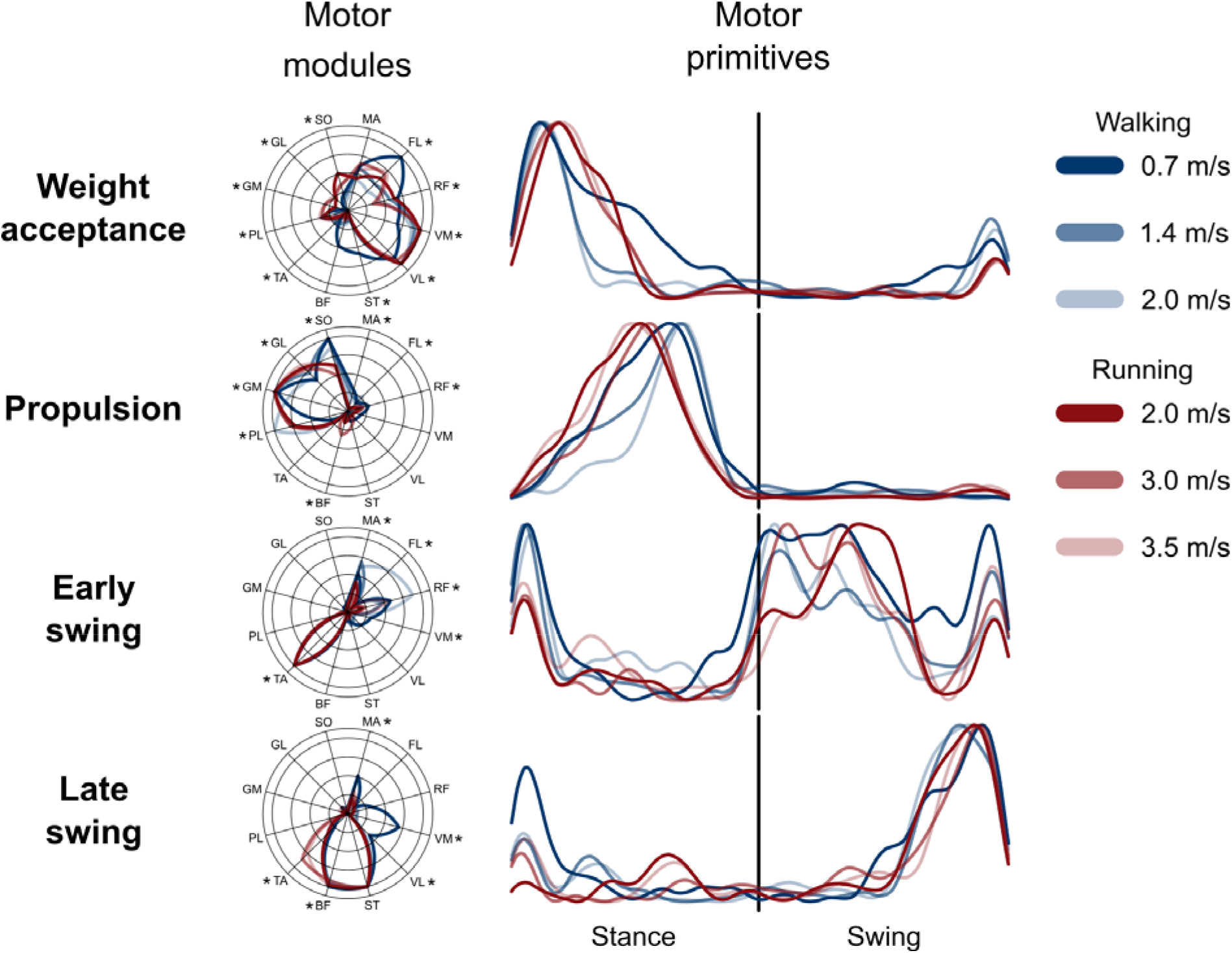
Muscle synergies for human walking and running at various speeds. Motor modules and motor primitives of the four fundamental synergies for human walking and submaximal running (average of all trials recorded in group 1). The motor modules are presented in polar coordinates on a normalized polar axis base. Each muscle contribution within one synergy can range from 0 to 1 (maximum radius length). Asterisks represent significant effect of speed (results of the *post-hoc* analysis, where relevant). For the motor primitives, the x-axis full scale represents the averaged gait cycle (with stance and swing normalized to the same amount of points and divided by a vertical line) and the y-axis the normalized amplitude. Muscle abbreviations: MA=*gluteus maximus*, FL=*tensor fasciæ latæ,* RF=*rectus femoris*, VM=*vastus medialis*, VL=*vastus lateralis*, ST=*semitendinosus*, BF=*biceps femoris*, TA=*tibialis anterior*, PL=*peroneus longus*, GM=*gastrocnemius medialis*, GL=*gastrocnemius lateralis*, SO=*soleus*.

**Figure 3.**
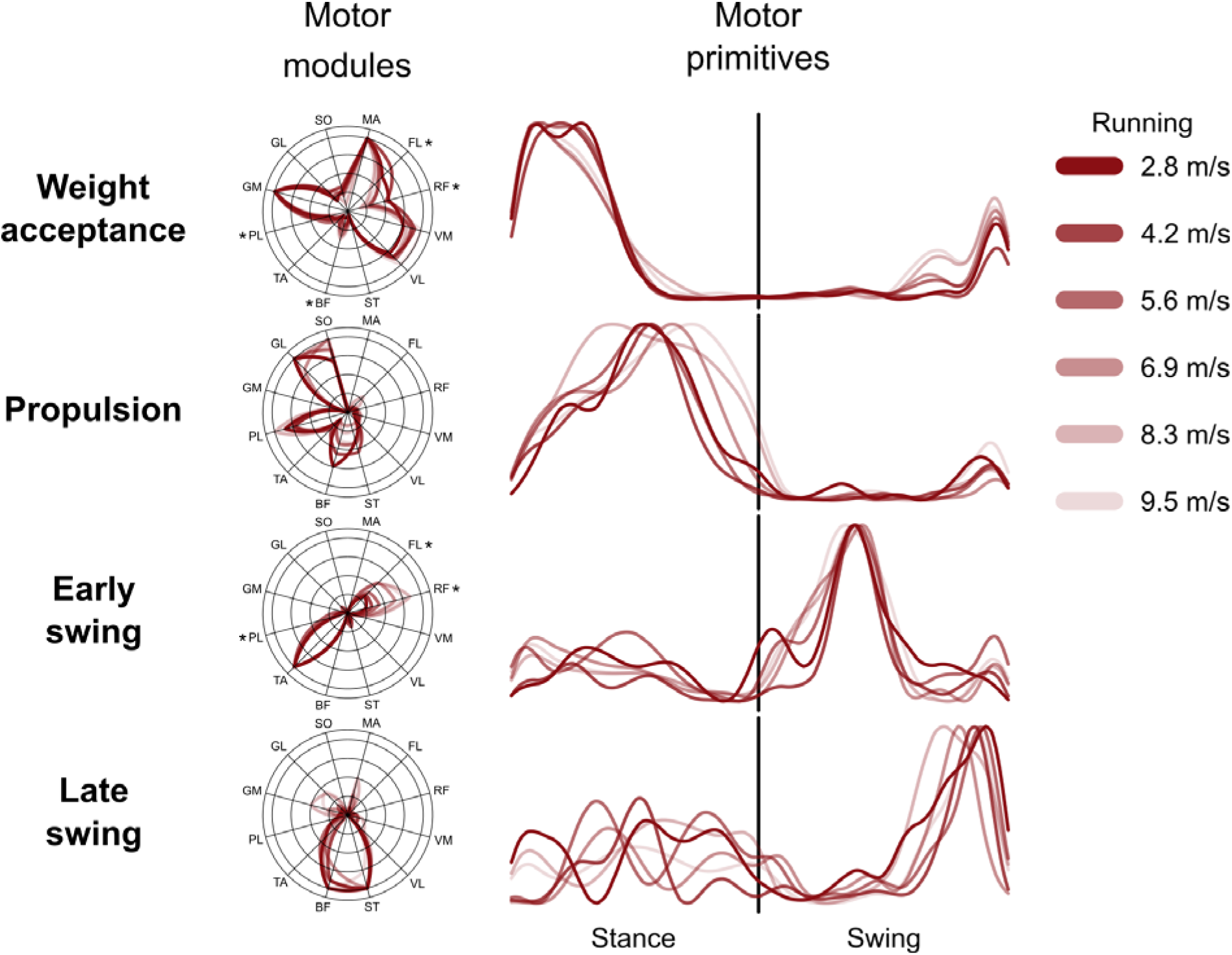
Muscle synergies for human running at various speeds. Motor modules and motor primitives of the four fundamental synergies for human submaximal and maximal running (average of all trials recorded in group 2). The motor modules are presented in polar coordinates on a normalized polar axis base. Each muscle contribution within one synergy can range from 0 to 1 (maximum radius length). Asterisks represent significant effect of speed (results of the *post-hoc* analysis, where relevant). For the motor primitives, the x-axis full scale represents the averaged gait cycle (with stance and swing normalized to the same amount of points and divided by a vertical line) and the y-axis the normalized amplitude. Muscle abbreviations: MA=*gluteus maximus*, FL=*tensor fasciæ latæ*, RF=*rectus femoris*, VM=*vastus medialis*, VL=*vastus lateralis*, ST=*semitendinosus*, BF=*biceps femoris*, TA=*tibialis anterior*, PL=*peroneus longus*, GM=*gastrocnemius medialis*, GL=*gastrocnemius lateralis*, SO=*soleus*.

**Table 1.** Frequency of occurrence of fundamental synergies. Even though the factorisation rank was not influenced by locomotion speed, not all the extracted synergies could be functionally classified as fundamental (i.e. not combined). This table reports the number of participants that showed the relevant fundamental synergies at each speed for both groups (G1 = walking and submaximal running, G2 = submaximal and maximal running).

| Group | Speed [m/s] |  | Synergy |  |  |  |
| --- | --- | --- | --- | --- | --- | --- |
|  |  |  | Weight acceptance | Propulsion | Early swing | Late swing |
| G1<br>(15 participants) | Walking | 0.7 | 10 | 15 | 13 | 6 |
|  |  | 1.4 | 13 | 15 | 9 | 15 |
|  |  | 2.0 | 13 | 15 | 7 | 15 |
|  | Running | 2.0 | 15 | 15 | 6 | 11 |
|  |  | 3.0 | 15 | 14 | 2 | 14 |
|  |  | 3.5 | 15 | 15 | 3 | 12 |
| G2 | Running | 2.8 | 15 | 13 | 8 | 4 |
| (15 participants) |  | 4.2 | 15 | 15 | 5 | 1 |
|  |  | 5.6 | 15 | 15 | 11 | 4 |
|  |  | 6.9 | 15 | 14 | 9 | 1 |
|  |  | 8.3 | 15 | 12 | 14 | 2 |
|  |  | 9.5 | 15 | 9 | 15 | 5 |

### Gait cycle parameters

An effect of speed (p < 0.001) was found in both groups for the cadence and the swing and stance times (Figure 4). When locomotion speed increased, cadence increased as well, while stance times decreased. In walking (G1), swing times decreased with increasing speed. In running (G1), swing times increased between 2.0 and 3.0 m/s, but were not significantly different at 3.0 and 3.5 m/s. In G2, swing times decreased with increasing speed after 4.2 m/s. The strike index during running was, in G1, of 0.23 ± 0.26 at 2.0 m/s, 0.22 ± 0.25 at 3.0 m/s and 0.24 ± 0.26 at 3.5 m/s, all indicating a rearfoot strike pattern. In G2, the strike index values during running were 0.50 ± 0.18 at 2.8 m/s, 0.57 ± 0.13 at 4.2 m/s, 0.62 ± 0.11 at 5.6 m/s, 0.65 ± 0.11 at 6.9 m/s, 0.70 ± 0.10 at 8.3 m/s and 0.74 ± 0.06 at 9.5 m/s, all indicating a mid/forefoot strike pattern.

**Figure 4.**
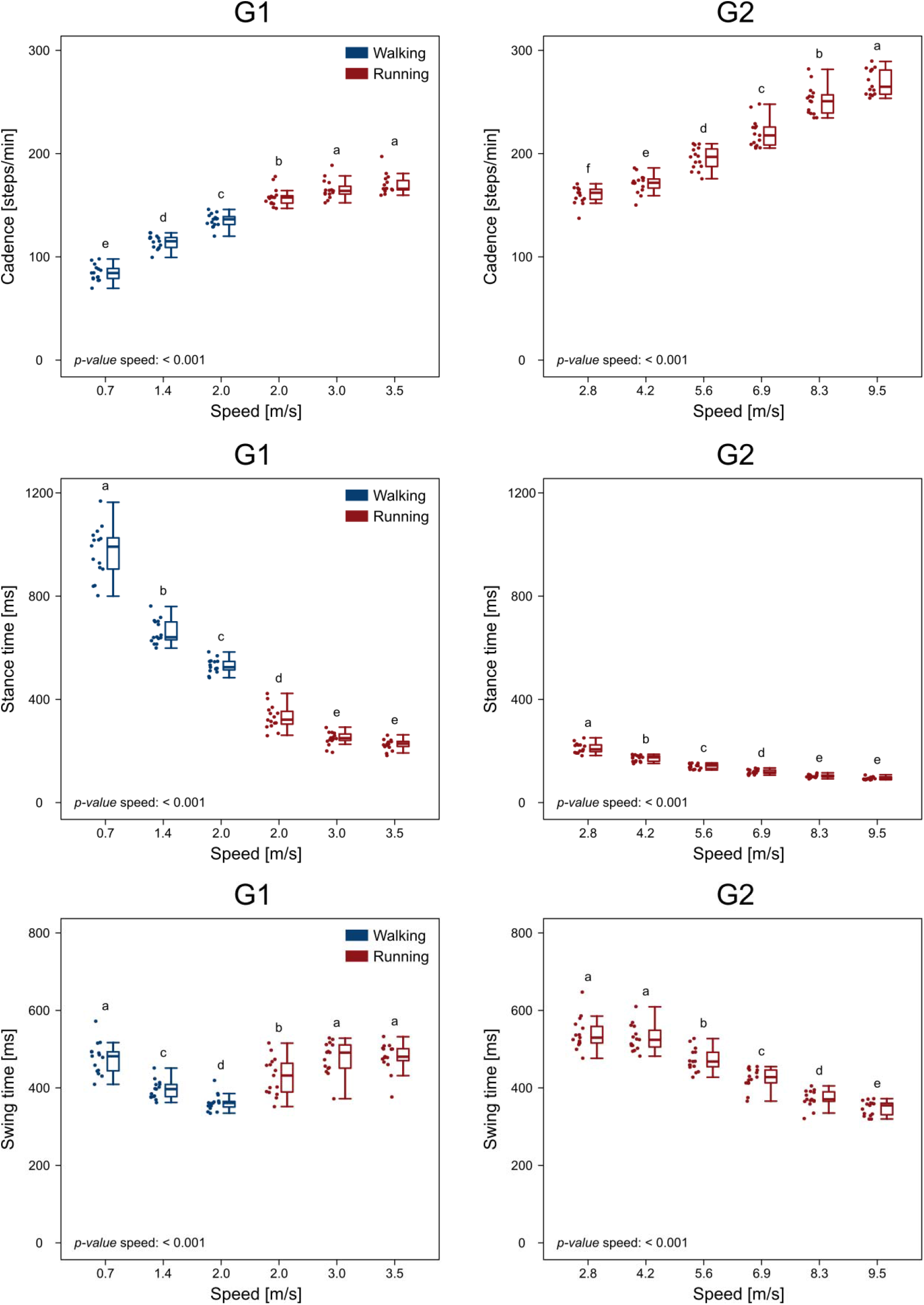
Gait cycle spatiotemporal parameters. Boxplots describing the cadence (in steps per minute), stance and swing times for the two groups (G1 = walking and submaximal running, G2 = submaximal and maximal running. Boxplots sharing the same letter (a, b, c, d, e) are not to be considered significantly different (results of the *post-hoc* analysis). Raw data points are presented to the left side of each boxplot.

### Higuchi’s fractal dimension of motor primitives

The HFD of motor primitives is reported in Figure 5. In both groups, the HFD was affected by speed (p < 0.001), with a global tendency towards a lower complexity (i.e. lower HFD) of motor primitives with increasing speed. Specifically, in G1 the highest complexity values were found in walking at 0.7 m/s, while walking at 1.4 and 2.0 m/s shared similar complexity of motor primitives; running from 2.0 to 3.5 m/s induced decreased HFD (Figure 5). In G2, the complexity decreased with speed until 8.3 m/s, while sprinting at 8.3 and 9.5 m/s did not produce any significant difference in the complexity of the motor primitives (Figure 5).

**Figure 5.**
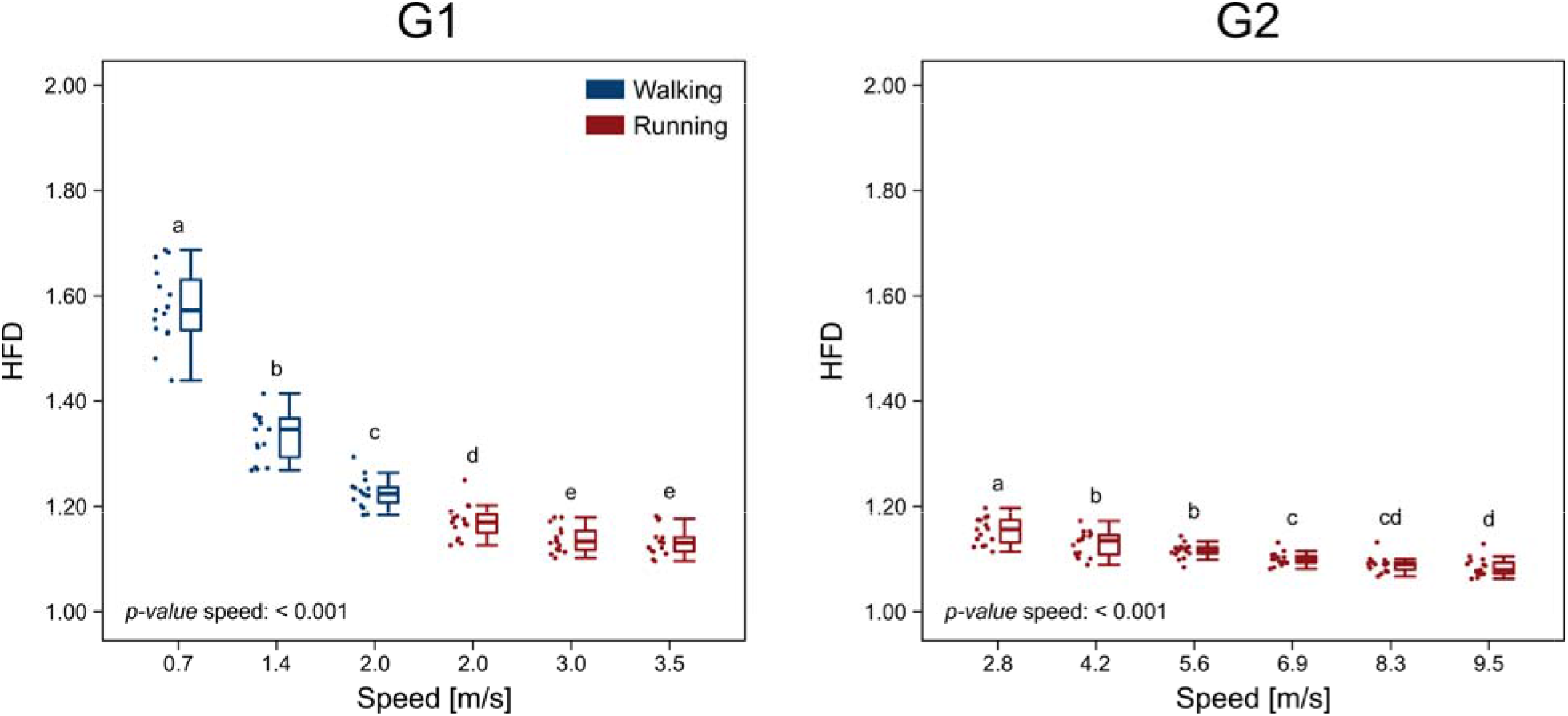
Higuchi’s fractal dimension of motor primitives. Boxplots describing the Higuchi’s fractal dimension (HFD) of the motor primitives extracted from the two groups (G1 = walking and submaximal running, G2 = submaximal and maximal running). Boxplots sharing the same letter (a, b, c, d, e) are not to be considered significantly different (results of the *post-hoc* analysis). Raw data points are presented to the left side of each boxplot.

### Width of motor primitives

The width of motor primitives, measured with the FWHM, was significantly affected by speed only for the primitives of the stance synergies (i.e. weight acceptance and propulsion) in both G1 and G2 (p < 0.001). The boxplots depicting the changes in FWHM with speed are shown in Figure 6. In G1, the weight acceptance primitives were wider in walking than in running, but speed did not play a role within the same locomotion type. The propulsion primitives, on the contrary, widened with increasing speed in both walking and running. In G2 there was a widening of the weight acceptance synergies after 5.6 m/s, while the propulsion synergies became wider with increasing speed at almost all speeds. The primitives of the early and late swing synergies did not show any change attributable to the different locomotion speed (early swing, G1: p = 0.133; late swing, G1: p = 0.029, *post-hoc* analysis did not confirm an effect of speed; early swing, G2: p = 0.385; late swing, G2: p = 0.391).

**Figure 6.**
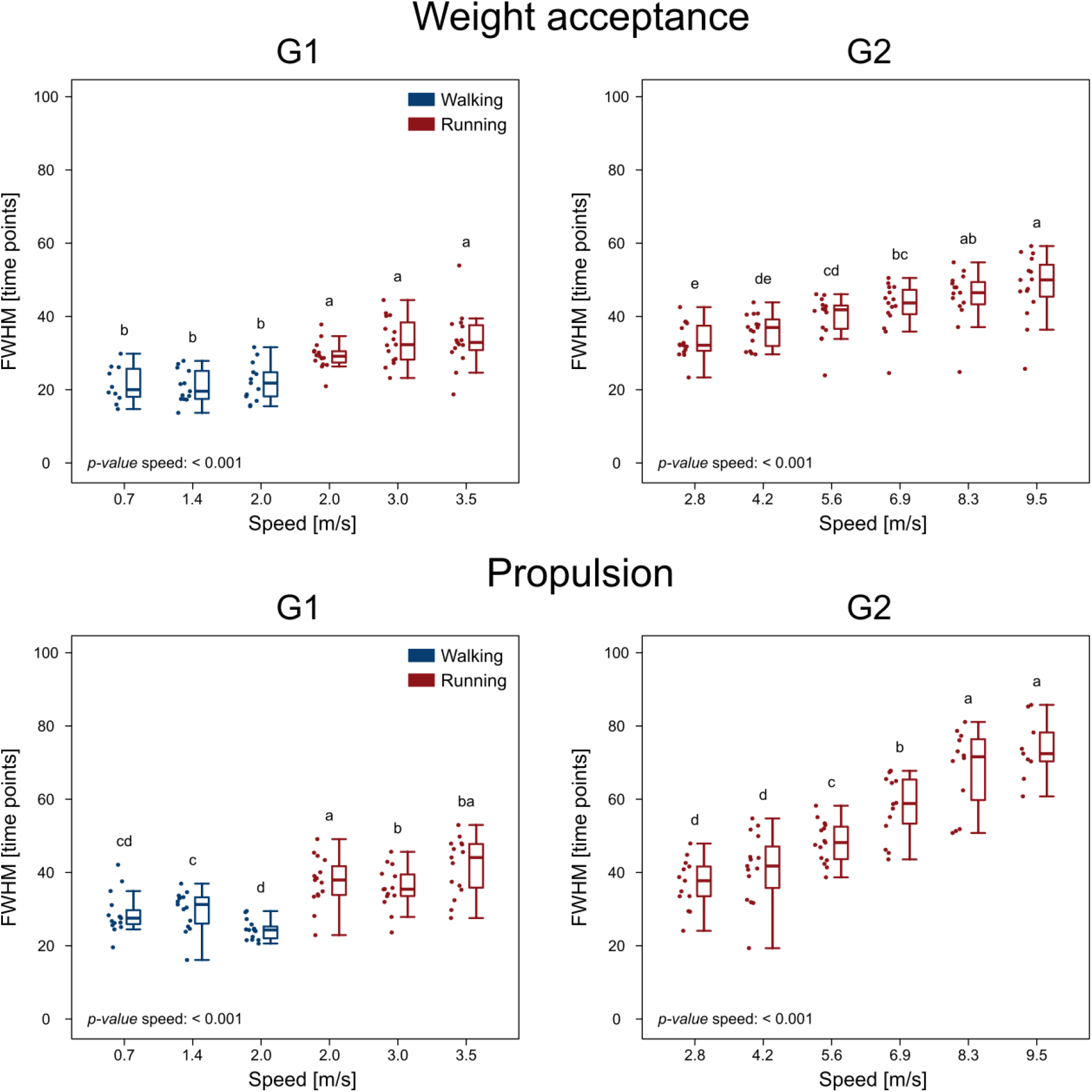
Width of motor primitives. Boxplots showing the width of motor primitives measured with the full width at half maximum (FWHM) in normalized time points for the two groups (G1 = walking and submaximal running, G2 = submaximal and maximal running). Only the primitives that showed significant effect of speed on the FWHM are showed (i.e. the motor primitives relative to the weight acceptance and propulsion synergies; the early and late swing synergies did not show any significant effect of speed on the FWHM of motor primitives). Boxplots sharing the same letter (a, b, c, d, e) are not to be considered significantly different (results of the *post-hoc* analysis). Raw data points are presented to the left side of each boxplot.

## Discussion

In this study, we described the modular organisation of muscle activations for human locomotion at various speeds. Our results showed that increasing the locomotion speed and transitioning from walking to running produced relatively longer (i.e. higher FWHM) and locally less complex (i.e. lower HFD) basic activation patterns (i.e. motor primitives). Moreover, we found a speed-dependent redistribution of muscle contributions (i.e. motor modules) in both walking and running. These findings provide evidence that muscle synergies must be spatially and temporally modulated to withstand the constraints imposed by high locomotion speeds.

### Longer, less complex motor primitives ensure robust control of high-speed locomotion

In the past, we used the FWHM of motor primitives as a measure of robustness [10,11,28]. Our conclusion was that wider (i.e. active for a relatively longer period of time) primitives indicate more robust control [10,11,28]. The overlap of chronologically-adjacent synergies increased the fuzziness [23,28,53] of temporal boundaries allowing for easier shifts between one synergy (or gait phase) and another [10,11]. Our conclusion fits the optimal feedback control theory, which postulates that motor systems selectively use feedback information to optimize an index of performance by combining sensory signals and motor commands [54–56]. For the CNS, this solution must come at the cost of reducing the accuracy or, as others called it, optimality [23] or efficiency [57]. Here, we found a clear widening of motor primitives at increasing locomotion speeds, but only in the synergies relevant for the stance phase (i.e. the weight acceptance and the propulsion synergies). This observation seems to confirm previous finding that more challenging locomotion conditions (in this case maximal as compared to submaximal running or fast as compared to slow walking) require more robust motor control achieved by widening the two motor primitives of the stance phase [10,11]. The FWHM did not only increase with running speed, but also when switching from walking to running. The challenge imposed by increased locomotion speed not only implies wider primitives, but lower complexity of neural control as well. Recently, we used the HFD, a measure of irregularity or complexity [20], to show that the motor primitives extracted from challenging locomotion conditions exhibit lower complexity than those associated with normal locomotion [11]. Specifically, we showed that older age and external perturbations produce lower complexity of motor primitives [11]. In this study, we found a similar behaviour depending on the speed at which our participants were walking or running. From the slowest (walking at 0.7 m/s) to the fastest speed (sprinting at 9.5 m/s), complexity decreased rather smoothly. In addition, as observed for the FWHM, primitives proved to be less complex in running than in walking, relative to the time-normalized gait cycle. This decrease in complexity can be interpreted as a strategy adopted by the CNS in parallel to the increased FWHM to robustly cope with the challenges imposed by high locomotion speeds. In fact, running allows less time for organising coordinated movements than walking [58] and a simplification of control could benefit robustness. Similarly, one could explain the need for lower complexity of motor primitives when locomotion speeds approach those of sprinting, with stance times well below 150 ms, swing times of less than 400 ms and cadence exceeding 250 steps per minute.

One possible reason for the increased FWHM when transitioning from walking to running could lie in the fact that the motor primitives for the two locomotion types have different shapes in the weight acceptance and propulsion synergy. Specifically, walking primitives are skewed to the left in the weight acceptance and to the right in the propulsion phase, while running primitives appear symmetrical with respect to the global maximum. In walking, the leading leg has a bigger angle at touchdown than in running [59]. The initial angle determines the position of the centre of mass, which reaches its highest (in walking) and lowest (in running) point in the middle of the stance phase [59]. This may physically constrain the production of forward forces in walking from the plantarflexors, which are major contributors in this phase [10,35,52,60], since only after half stance there can be propulsion, while in running it can happen earlier [2]. This could explain the physiological constraint to the recruitment timing of those muscles responsible for the weight acceptance (mostly knee extensors) and propulsion (mostly plantarflexors) phase. Additionally, the question remains as to why HFD values are lower when the walking or running speed increases and in walking compared to running. Due to its definition (EQ4), the HFD depends on the signal-to-noise ratio [61], but some precautions can be taken to reduce the influence of the signal-to-noise ratio on the outcomes (i.e. minimum subtraction and normalisation to the maximum). Nevertheless, due to the summation term in EQ4, which represents the absolute value of the successive differences of each motor primitive’s ordinates, calculated with lag k (see methods), curves with bigger FWHM will have a lower L(k). From a physiological point of view this could mean that the CNS deals with the challenge of controlling locomotion at high speeds by increasing the FWHM of control signals, a solution that results in locally less complex motor primitives (i.e. lower HFD) relative to the time-normalized gait cycle. Lower complexity indicates lower nonlinearity of the physiological signal [45], in our case the motor primitives. It has been shown that the complexity of electroencephalographic activity is reduced by degeneration and dysfunction of neural networks, e.g. due to aging, neurodegenerative diseases, brain injury and stroke [45]. As mentioned before, the complexity of motor primitives for locomotion decreases with aging and when external perturbations are introduced [11]. Following these observations, it is tempting to associate decreased neural complexity with internally or externally imposed constraints to movement. The ability to modulate complexity might in fact be a determinant of sprinting performance. Based on the present findings, it could be possible to speculate that interventions focused on the regulation of motor primitive complexity could be used to assess and possibly improve the performance high-speed locomotion (walking and running). This idea might path the way for the establishment of future training intervention protocols aimed not only at athletes but possibly at patients suffering from neurological diseases as well. However, not only the timing is important for coordinated, robust control of fast locomotion, but the recruitment of the appropriate muscle groups is of critical importance as well.

### Fast locomotion requires different muscle contributions

Our results also show that biarticular muscles have a speed-dependent function in both walking and running. The RF as a biarticular muscle (i.e. crossing hip and knee) has the double function of extending the knee and flexing the hip [62]. In our analysis, it is possible to notice that when increasing both the walking and running speed, the relative contribution of the RF shifts from the weight acceptance synergy (typical domain of the other two quadriceps muscle recorded, the VM and VL) to the early swing synergy. This might be an indication of the double function of this muscle: knee extensor and hip flexor. In fact, in the weight acceptance phase of the gait cycle, the quadriceps muscle group is known to aid the deceleration and support of the body mass, acting in a quasi-isometric way at the muscle fascicle level [63]. In the early swing phase, the *iliopsoas (iliacus* and *psoas)* muscle group is one of the major contributors to the hip flexion, which starts right after the lift-off and continues for around three quarters of the swing phase [64–68]. At high walking and running speeds, the RF shifts its contribution from the weight acceptance synergy (working as knee extensor) to the early swing synergy (working as hip flexor [69,70] and as knee extensor to counteract the inertial flexion of the knee joint in the last third of the swing phase [66,67]). Our results show that this behaviour is achieved by modulating the relative contribution to the motor modules of the two relevant synergies. A similar behaviour is evident in the relative contribution of the FL muscle to the motor modules of the weight acceptance and early swing synergies, in both walking and running.

Another outcome of this study was the different contribution of the GM muscle for the participants of G1 and G2. In G1, as found in the past [10,33,35,38,52], the GM and GL were crucial contributors to the propulsion synergy, together with the SO and PL. In G2, though, the contribution of the GM shifted to the weight acceptance phase at both maximal and submaximal speeds. During running, the participants included in G1 (recreational long-distance runners) adopted a different landing preparation strategy than those of G2 (national level sprinters) even at similar speeds. This is visible in the relative contribution of the TA muscle to the motor modules of the late swing synergy. It has been shown that the foot strike pattern is intimately linked to the modular organisation of this synergy, since rearfoot strikers need to dorsiflex the foot more than mid/forefoot strikers before touchdown [38]. In the present data, the TA contribution to the late swing synergy was considerable in G1 (average motor module value excluding running at 2.0 m/s: 0.57 ± 0.39, the minimum being 0 and the maximum 1), but reduced in G2 (average motor module value: 0.29 ± 0.30). This observation is confirmed by the fact that, on average, the participants of G1 had a rearfoot strike pattern, contrary to the mid/forefoot strike pattern of the participants included in G2 (see results). The different kinematics of the ankle joint at touchdown might be a reason for the different contribution of the GM and GL muscles to the weight acceptance and propulsion synergies in G1 and G2. Similarly, it is evident in G2, compared to G1, an extended contribution of the ST and BF to the propulsion phase. In G2, the two biarticular hamstrings are used both in the propulsion synergy (likely mostly as hip extensors) and in the late swing synergy, a feature that is not visible in G1, where the ST and BF are exclusively contributing to the late swing.

### Conclusion

Our results show that wider, less complex muscle activation patterns are needed to cope with the challenges imposed by increased locomotion speeds. The width, complexity and modularity of muscle synergies can be regulated to ensure robust locomotion control even at very high speeds. This stands for both walking and running, with running showing generally less complex, wider motor primitives than walking.

## Materials and Methods

This study was reviewed and approved by the Ethics Committees of the Humboldt-Universität zu Berlin and Osaka University of Health and Sport Sciences. All the participants gave written informed consent for the experimental procedure, in accordance with the Declaration of Helsinki. Experimental protocols

For the two experimental protocols we recruited 30 healthy male volunteers and divided them into two groups. The first group of 15 recreational long-distance runners (henceforth G1, height 178 ± 6 cm, body mass 71 ± 6 kg, age 33 ± 6 years, 43 ± 21 km/week running volume, personal best mark over 10 km 37.4 ± 3.2 minutes, means ± standard deviation) was assigned to the first experimental protocol conducted at the Humboldt-Universität zu Berlin (Germany). The second group of 15 sprint athletes (G2, height 172 ± 4 cm, body mass 65 ± 3 kg, age 21 ± 2 years, personal best mark over 100 m 10.74 ± 0.23 s) was assigned to the second experimental protocol conducted at the Osaka University of Health and Sport Sciences (Japan). All the participants completed a self-selected warm-up running on a treadmill, typically lasting between 3 and 5 min [33,34]. After being instructed about the protocol, they completed a different set of measurements, depending on the protocol they were assigned to.

The experimental protocol of G1 consisted of walking (at 0.7, 1.4, and 2.0 m/s) and submaximal running (at 2.0, 3.0, and 3.5 m/s) on a single-belt treadmill (mercury, H-p-cosmos Sports & Medical GmbH, Nussdorf, Germany) equipped with a pressure plate recording the plantar pressure distribution at 120 Hz (FDM-THM-S, zebris Medical GmbH, Isny im Allgäu, Germany). The speeds were chosen as follows: walking at 1.4 m/s and running at 3.0 m/s are the commonly reported average comfortable locomotion speeds [34,35]; 2.0 m/s is the typical walk-to-run transition speed [36,37]; the other two speeds were chosen to extend the range of investigation.

The experimental protocol of G2 consisted of running (at 2.8, 4.2, 5.6, 6.9, 8.3, and 9.5 m/s) on a single-belt treadmill (Fully Instrumented Treadmill, Bertec co., Columbus, OH, USA) modified to reach the maximum speed of 9.5 m/s and equipped with force sensors to record the 3D ground reaction forces at 1 kHz. The highest sprinting speed was chosen to match the average pace used by the participants to run 100 m close to their personal best time.

### EMG recordings

The muscle activity of the following 12 ipsilateral (right side) muscles was recorded in both groups: *gluteus maximus* (MA), *tensor fasciæ latæ* (FL), *rectus femoris* (RF), *vastus medialis* (VM), *vastus lateralis* (VL), *semitendinosus* (ST), *biceps femoris* (long head, BF), *tibialis anterior* (TA), *peroneus longus* (PL), *gastrocnemius medialis* (GM), *gastrocnemius lateralis* (GL) and *soleus* (SO). The electrodes were positioned as extensively reported previously [19,33]. After around 60 s habituation [10] in G1 or after a mild acceleration of the belt in G2 (lasting 5 to 10 s depending on the speed), we recorded one trial for each participant with an acquisition frequency of 2 kHz by means of a 16-channel wireless bipolar EMG system (Wave Plus wireless EMG, Cometa srl, Bareggio, Italy). For the EMG recordings, we used foam-hydrogel electrodes with snap connector (H124SG, Medtronic plc, Dublin, Ireland). The first 30 gait cycles of the recorded trial were considered for subsequent analysis [33]. Exceptions (13 out of 15 participants of G2) to this rule were applied if the participants could not sustain the imposed speed for a sufficient number of gait cycles (an event occurring only at the higher sprinting speed). All the recordings can be downloaded from the supplementary data set, which is accessible at Zenodo (DOI: 10.5281/zenodo.3764761).

### Gait cycle parameters

The gait cycle breakdown (foot touchdown and lift-off timing) was obtained by the elaboration of the data acquired by the pressure (G1) and force (G2) plates with validated algorithms that were reported previously [34]. Other calculated gait spatiotemporal parameters were: cadence (i.e. number of steps per minute), stance and swing times and the strike index, calculated as the distance from the heel to the centre of pressure at impact relative to total foot length [34]. Strike index values range from 0 to 1, denoting the most posterior and the most anterior point of the shoe, respectively [38]. Values from 0.00 to 0.33 are indication of a rearfoot strike pattern, while values from 0.34 to 1.00 represent a mid/forefoot strike pattern [34].

### Muscle synergies extraction

Muscle synergies data were extracted from the recorded EMG activity through a custom script (R v3.6.3, R Core Team, 2020, R Foundation for Statistical Computing, Vienna, Austria) using the classical Gaussian non-negative matrix factorisation (NMF) algorithm [10,33,35,39]. The raw EMG signals were band-pass filtered within the acquisition device (cut-off frequencies 10 and 500 Hz). Then the signals were high-pass filtered, full-wave rectified and lastly low-pass filtered using a 4^th^ order IIR Butterworth zero-phase filter with cut-off frequencies 50 Hz (high-pass) and 20 Hz (low-pass for creating the linear envelope of the signal) as previously described [10]. After subtracting the minimum, the amplitude of the EMG recordings obtained from the single trials was normalized to the maximum activation recorded for every individual muscle (i.e. every EMG channel was normalized to its maximum in every trial) [19,33]. Each gait cycle was then time-normalized to 200 points, assigning 100 points to the stance and 100 points to the swing phase [10,19,33,38]. The reason for this choice is twofold [33]. First, dividing the gait cycle into two macro-phases helps the reader understanding the temporal contribution of the different synergies, diversifying between stance and swing. Second, normalising the duration of stance and swing to the same number of points for all participants (and for all the recorded gait cycles of each participant) makes the interpretation of the results independent from the absolute duration of the gait events. Synergies were then extracted through NMF as previously described [10,33]. The 12 muscles listed above were considered for the analysis, (MA, FL, RF, VM, VL, ST, BF, TA, PL, GM, GL and SO). The m = 12 time-dependent muscle activity vectors were grouped in a matrix V with dimensions m × n (m rows and n columns). The dimension n represented the number of normalized time points (i.e. 200*number of gait cycles). The matrix V was factorized using NMF so that V ≈ V_R_ = WH. The new matrix V_R_, reconstructed multiplying the two matrices W and H, approximates the original matrix V. The motor primitives [35,40] matrix H contained the time-dependent coefficients of the factorisation with dimensions r × n, where the number of rows r represents the minimum number of synergies necessary to satisfactorily reconstruct the original set of signals V. The motor modules [35,41] matrix W, with dimensions m × r, contained the time-invariant muscle weightings, which describe the relative contribution of single muscles within a specific synergy (a weight was assigned to each muscle for every synergy). H and W described the synergies necessary to accomplish the required task (i.e. walking or swimming). The update rules for W and H are presented in Equation (EQ1) and Equation (EQ2).

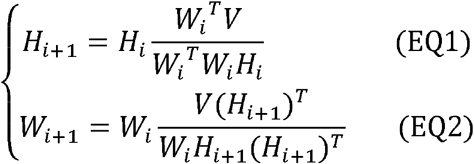

The quality of reconstruction was assessed by measuring the coefficient of determination R^2^ between the original and the reconstructed data (V and V_R_, respectively). The limit of convergence for each synergy was reached when a change in the calculated R^2^ was smaller than the 0.01% in the last 20 iterations [35] meaning that, with that amount of synergies, the signal could not be reconstructed any better. This operation was first completed by setting the number of synergies to 1. Then, it was repeated by increasing the number of synergies each time, until a maximum of 9 synergies. The number 9 was chosen to be lower than the number of muscles, since extracting a number of synergies equal to the number of measured EMG activities would not reduce the dimensionality of the data. Specifically, 9 is the rounded 75% of 12, which is the number of considered muscles [19]. For each synergy, the factorisation was repeated 10 times, each time creating new randomized initial matrices W and H, in order to avoid local minima [42]. The solution with the highest R^2^ was then selected for each of the 9 synergies. To choose the minimum number of synergies required to represent the original signals, the curve of R^2^ values versus synergies was fitted using a simple linear regression model, using all 9 synergies. The mean squared error [43] between the curve and the linear interpolation was then calculated. Afterwards, the first point in the R^2^-vs.-synergies curve was removed and the error between this new curve and its new linear interpolation was calculated. The operation was repeated until only two points were left on the curve or until the mean squared error fell below 10^-4^. This was done to search for the most linear part of the R^2^-versus-synergies curve, assuming that in this section the reconstruction quality could not increase considerably when adding more synergies to the model.

### Higuchi’s fractal dimension of motor primitives

To assess the local complexity [44] of motor primitives, we calculated the Higuchi’s fractal dimension (HFD), assuming that these time series exhibit self-affinity properties [11,20,21,45–48]. Following the procedure first described by Higuchi[20], for each motor primitive *H(t)*[*H(1), H(2)*, … *H(n)*], *k* sets of new time series must be constructed, where *k* is an integer interval time and 2 < *k* < *k_max_*:

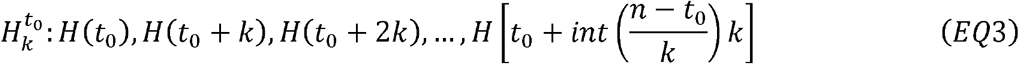

where *t*_0_ is the first sample at initial time. The non-Euclidean length of each curve was defined as

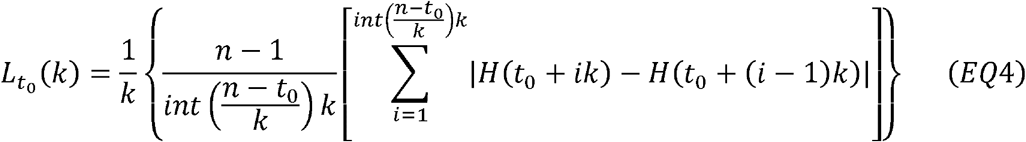

and for every considered *k* step the length of the motor primitive was defined as the average of the *k* sets of lengths as

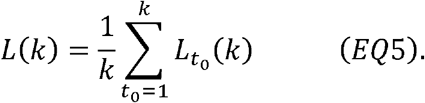

If L(k) ∝ *k^*HFD^*, then the curve is fractal with dimension HFD and this should lead the plot of log(L(k)) versus log(1/k) to fall on a straight line with slope-HFD. The values of the HFD range from 1 (e.g. for a smooth linear time series) to 2 (e.g. for random white noise) and are independent on the amplitude of the signal, since the curve log(L(k)) versus log(1/k) changes intercept but not slope if the same signal is multiplied or divided [45,47]. For each trial, the HFD of the primitives obtained by NMF was calculated separately and then averaged, so that each trial ultimately consisted of one HFD value [11]. Following suggestions from previous studies, *k_max_* was chosen as the most linear part of the log-log plot, which in our data led us to choose *k_max_* = 10 [48,49].

### Width of motor primitives

We compared motor primitives by evaluating the full width at half maximum (FWHM), a metric useful to describe the duration of activation patterns [3,10,19,25]. The FWHM was calculated cycle-by-cycle after subtracting the cycle’s minimum as the number of points exceeding each cycle’s half maximum, and then averaged [25]. The FWHM (and just this parameter) was calculated only for the motor primitives relative to fundamental synergies. A fundamental synergy can be defined as an activation pattern whose motor primitive shows a single main peak of activation [10]. When two or more fundamental synergies are blended into one, a combined synergy appears. Combined synergies usually constitute, in our locomotion data, 10 to 30% of the total extracted synergies. While fundamental synergies can be compared given their similar function (i.e. motor primitives and motor modules are comparable since they serve a specific task within the step cycle), combined synergies often differ from one another making their classification impossible. Due to the lack of consent in the literature on how to interpret them, we excluded the combined synergies from the FWHM (but not the HFD) analysis.

### Functional classification of muscle synergies

The recognition of fundamental synergies was carried out by clustering similar motor primitives through NMF, using the same algorithm employed for synergy extraction with the maximum number of synergies set to the maximum factorisation rank plus one. The obtained “principal shapes” (four for G1 walking, G1 running and G2 running) were then compared to the motor primitives in order to cluster similar shapes. A primitive was considered similar to one of the principal shapes if the NMF weight was equal at least to the average of all weights. Of all the primitives that satisfied this condition, we then calculated the R^2^ with the relevant principal shape. If the R^2^ was at least the 25% (or four times if the R^2^ was negative) of the average R^2^ obtained by comparing all the remaining primitives with their own principal shape, we confirmed the synergy as fundamental and classified it based on function. Primitives that were not clustered, were labelled as combined.

### Statistics

To investigate the effect of locomotion speed on the factorisation rank, gait parameters, HFD and FWHM of motor primitives, and motor modules we fitted the data using a generalized linear model with Gaussian error distribution. The homogeneity of variances was tested using the Levene’s test. If the residuals were normally distributed, we carried out a one-way repeated measures ANOVA with type II sum of squares for the dependent variables factorisation rank, cadence, stance and swing time, HFD and FWHM, the independent variable being the locomotion speed. If the normality assumptions on the residuals were not met, we used the non-parametric Kruskal-Wallis test. For the motor modules, we carried out a two-way repeated measures ANOVA with type II sum of squares, the independent variables being the speed and the muscles. If the normality assumptions on the residuals were not met, we used a robust (rank-based) ANOVA from the R package Rfit (function “raov”) [50,51]. We then performed a least significant difference *post-hoc* analysis with false discovery rate adjustment of the *p*-values. All the significance levels were set to α = 0.05 and the statistical analyses were conducted using R v3.6.3.

### Data availability

In the supplementary data set accessible at Zenodo (DOI: 10.5281/zenodo.3764761) we made available: a) the metadata with anonymized participant information, b) the raw EMG, c) the touchdown and lift-off timings of the recorded limb, d) the filtered and time-normalized EMG, e) the muscle synergies extracted via NMF and f) the code to process the data, including the scripts to calculate the HFD of motor primitives. In total, 180 trials from 30 participants are included in the supplementary data set.

The file “metadata.dat” is available in ASCII and RData format and contains:

- Code: the participant’s code
- Group: the experimental group in which the participant was involved (G1 = walking and submaximal running; G2 = submaximal and maximal running)
- Sex: the participant’s sex (M or F)
- Speeds: the type of locomotion (W for walking or R for running) and speed at which the recordings were conducted in 10*[m/s]
- Age: the participant’s age in years
- Height: the participant’s height in [cm]
- Mass: the participant’s body mass in [kg]
- PB: 100 m-personal best time (for G2).

The files containing the gait cycle breakdown are available in RData format, in the file named “CYCLE_TIMES.RData”. The files are structured as data frames with as many rows as the available number of gait cycles and two columns. The first column named “touchdown” contains the touchdown incremental times in seconds. The second column named “stance” contains the duration of each stance phase of the right foot in seconds. Each trial is saved as an element of a single R list. Trials are named like “CYCLE_TIMES_P20_R_20,” where the characters “CYCLE_TIMES” indicate that the trial contains the gait cycle breakdown times, the characters “P20” indicate the participant number (in this example the 20th), the character “R” indicate the locomotion type (W=walking, R=running), and the numbers “20” indicate the locomotion speed in 10*m/s (in this case the speed is 2.0 m/s). Please note that the following trials include less than 30 gait cycles (the actual number shown between parentheses): P16_R_83 (20), P16_R_95 (25), P17_R_28 (28), P17_R_83 (24), P17_R_95 (13), P18_R_95 (23), P19_R_95 (18), P20_R_28 (25), P20_R_42 (27), P20_R_95 (25), P22_R_28 (23), P23_R_28(29), P24_R_28 (28), P24_R_42 (29), P25_R_28 (29), P25_R_95 (28), P26_R_28 (29), P26_R_95 (28), P27_R_28 (28), P27_R_42 (29), P27_R_95 (24), P28_R_28 (29), P29_R_95 (17).

The files containing the raw, filtered and the normalized EMG data are available in RData format, in the files named “RAW_EMG.RData” and “FILT_EMG.RData”. The raw EMG files are structured as data frames with as many rows as the amount of recorded data points and 13 columns. The first column named “time” contains the incremental time in seconds. The remaining 12 columns contain the raw EMG data, named with muscle abbreviations that follow those reported above. Each trial is saved as an element of a single R list. Trials are named like “RAW_EMG_P03_R_30”, where the characters “RAW_EMG” indicate that the trial contains raw emg data, the characters “P03” indicate the participant number (in this example the 3rd), the character “R” indicate the locomotion type (see above), and the numbers “30” indicate the locomotion speed (see above). The filtered and time-normalized emg data is named, following the same rules, like “FILT_EMG_P03_R_30”.

The files containing the muscle synergies extracted from the filtered and normalized EMG data are available in RData format, in the files named “SYNS_H.RData” and “SYNS_W.RData”. The muscle synergies files are divided in motor primitives and motor modules and are presented as direct output of the factorisation and not in any functional order. Motor primitives are data frames with 6000 rows and a number of columns equal to the number of synergies (which might differ from trial to trial) plus one. The rows contain the time-dependent coefficients (motor primitives), one column for each synergy plus the time points (columns are named e.g. “time, Syn1, Syn2, Syn3”, where “Syn” is the abbreviation for “synergy”). Each gait cycle contains 200 data points, 100 for the stance and 100 for the swing phase which, multiplied by the 30 recorded cycles, result in 6000 data points distributed in as many rows. This output is transposed as compared to the one discussed in the methods section to improve user readability. Each set of motor primitives is saved as an element of a single R list. Trials are named like “SYNS_H_P12_W_07”, where the characters “SYNS_H” indicate that the trial contains motor primitive data, the characters “P12” indicate the participant number (in this example the 12th), the character “W” indicate the locomotion type (see above), and the numbers “07” indicate the speed (see above). Motor modules are data frames with 12 rows (number of recorded muscles) and a number of columns equal to the number of synergies (which might differ from trial to trial). The rows, named with muscle abbreviations that follow those reported above, contain the time-independent coefficients (motor modules), one for each synergy and for each muscle. Each set of motor modules relative to one synergy is saved as an element of a single R list. Trials are named like “SYNS_W_P22_R_20”, where the characters “SYNS_W” indicate that the trial contains motor module data, the characters “P22” indicate the participant number (in this example the 22nd), the character “W” indicates the locomotion type (see above), and the numbers “20” indicate the speed (see above). Given the nature of the NMF algorithm for the extraction of muscle synergies, the supplementary data set might show non-significant differences as compared to the one used for obtaining the results of this paper.

The files containing the HFD calculated from motor primitives are available in RData format, in the file named “HFD.RData”. HFD results are presented in a list of lists containing, for each trial, 1) the HFD, and 2) the interval time *k* used for the calculations. HFDs are presented as one number (mean HFD of the primitives for that trial), as are the interval times *k*. Trials are named like “HFD_P01_R_95”, where the characters “HFD” indicate that the trial contains HFD data, the characters “P01” indicate the participant number (in this example the 1st), the character “R” indicates the locomotion type (see above), and the numbers “95” indicate the speed (see above). All the code used for the pre-processing of EMG data, the extraction of muscle synergies and the calculation of HFD is available in R format. Explanatory comments are profusely present throughout the script “muscle_synergies.R”.

## Acknowledgments

The authors are grateful to all the participants that showed great commitment and interest during the experiments. The authors disclose any professional relationship with companies or manufacturers who might benefit from the results of the present study.

## Author contributions

Conceptualisation: A.S., M.I., and A.A.; Data curation: A.S., and Y.K.; Formal analysis: A.S., and Y.K.; Investigation: A.S., A.E., Y.K., K.K., and M.I.; Methodology: A.S., M.I., and A.A.; Project administration: A.S., M.I., and A.A.; Resources: A.S., A.E., Y.K., K.K., M.I., and A.A.; Software: A.S.; Supervision: A.S., M.I., and A.A.; Validation: A.S., A.E., Y.K., K.K., M.I., and A.A.; Visualisation: A.S.; Writing – original draft: A.S., and A.A.; Writing – review & editing: A.S., A.E., Y.K., K.K., M.I., and A.A.

